# Oral administration of Angiotensin-(1–7) included in hydroxypropyl β-cyclodextrin attenuates cardiac hypertrophy and inflammation in a mouse model of Takotsubo cardiomyopathy

**DOI:** 10.64898/2026.10.01.756123

**Authors:** Maria Luiza-Dias, Bruno Sanches, Julia Rezende-Ribeiro, Vladimir Gorshkov, Frank Kjeldsen, Silvia Guatimosim, Cibele Rocha-Resende, Thiago Verano-Braga

**Affiliations:** Departamento de Fisiologia e Biofísica, Instituto de Ciências Biológicas, Universidade Federal de Minas Gerais, Antônio Carlos 6627, 31270-901 Belo Horizonte, Minas Gerais, Brazil; Department of Biochemistry and Molecular Biology, University of Southern Denmark, Campusvej 55, 5230 Odense M, Denmark

**Keywords:** Takotsubo cardiomyopathy, Angiotensin-(1–7), Renin-angiotensin system Isoproterenol, Cardiac hypertrophy, Cardiac inflammation

## Abstract

**Background:** Takotsubo cardiomyopathy (TTC) is a stress-induced cardiomyopathy characterized by acute cardiac dysfunction, inflammation, and hypertrophy, yet no therapy directly targets the underlying myocardial injury. Clinical studies have reported elevated circulating Angiotensin II (Ang II) levels in TTC patients, suggesting dysregulation of the renin-angiotensin system (RAS). Given the established cardioprotective effects of Angiotensin-(1-7) [Ang-(1–7)] in other cardiovascular diseases, we hypothesized that oral administration of Ang-(1–7) could attenuate acute cardiac remodeling during TTC.

**Methods:** Female C57BL/6J mice were subjected to a single high-dose isoproterenol (ISO) injection (300 mg/Kg, intraperitoneal) to induce TTC and received daily oral treatment with Ang-(1-7) complexed with hydroxypropyl β-cyclodextrin [Ang-(1-7)-CD] or vehicle (CD), for 4 days, beginning one day before ISO administration. Cardiac proteome alterations were characterized by tandem mass tag (TMT)-based quantitative proteomics. Cardiac hypertrophy was assessed by heart weight normalized to tibial length and cardiomyocyte cross-sectional area. Myocardial inflammation was evaluated by histopathology, CD68 immunostaining, and flow cytometric analysis of infiltrating leukocyte populations.

**Results:** We observed a distinct cardiac proteome signature in Ang-(1-7)-treated ISO mice, predicting inhibition of pathways associated with cardiac hypertrophy and adverse remodeling. These molecular changes were accompanied by reduced heart weight normalized to tibial length and smaller cardiomyocyte cross-sectional area compared with vehicle-treated ISO mice. Oral Ang-(1-7) also attenuated myocardial inflammation, as evidenced by lower inflammatory scores, reduced CD68⁺ macrophage infiltration, decreased numbers of CD45⁺ leukocytes, and marked suppression of Ly6G⁺ neutrophil and CCR2⁺ monocyte recruitment to the heart. These protective effects were associated with improved survival following ISO challenge. MS data are available via ProteomeXchange (PXD082872).

**Conclusions:** Oral administration of Ang-(1–7) attenuated acute cardiac hypertrophy and inflammation in experimental TTC while inducing cardiac proteome changes consistent with reduced adverse remodeling. These findings identify activation of the protective RAS axis as a promising therapeutic strategy for limiting myocardial injury during TTC.

## INTRODUCTION

Takotsubo cardiomyopathy (TTC) was first described in the 1990s ^1^, and is also referred to as stress-induced cardiomyopathy. TTC is a cardiovascular disorder that predominantly affects women and is typically triggered by intense physical or emotional stress, including substance abuse, accidents, environmental disasters, or bereavement ^2^. Accumulating evidence suggests that intense stressors induce excessive release of catecholamines, leading to supraphysiological stimulation of cardiac β-adrenergic receptors (β-ARs) and maladaptive cardiac remodeling ^3^. TCC patients often present ST-segment elevation on electrocardiography, chest pain, dyspnea, and characteristic apical ballooning of the left ventricle (LV) observed by echocardiography ^4^. Importantly, the clinical manifestation of TTC closely resembles that of acute myocardial infarction, frequently leading to diagnostic confusion and misclassification. However, TTC can be distinguished from myocardial infarction by markedly elevated plasma concentrations of epinephrine and norepinephrine, the absence of obstructive coronary artery disease, and the presence of reversible LV apical ballooning ^1^. Although TTC has long been considered a condition with a low mortality rate, with generally favorable prognosis, recent studies have demonstrated persistent maladaptive structural, functional, and inflammatory alterations in the myocardium of patients even after apparent clinical recovery ^5^, highlighting the underestimated severity of this cardiomyopathy. Although TTC is now more readily recognized, its historical misdiagnosis as acute myocardial infarction delayed the development of targeted therapies. Consequently, current treatment remains focused on supportive care and the management of complications rather than on interventions that directly address the underlying mechanisms of myocardial injury, underscoring the need for mechanistically driven therapeutic approaches.

Clinical studies have demonstrated that TTC patients have elevated levels of serum angiotensin II (Ang II) ^6^, a peptide of the renin-angiotensin system (RAS) canonic axis ^7,8^. When Ang II binds to AT1 receptor (AT1R), it promotes aldosterone secretion from the adrenal cortex, increases heart rate and vascular tone (vasoconstriction), and enhances sympathetic nervous system activity, resulting in elevated blood pressure ^8^. However, the hyperactivation of the Ang II/AT1R axis is related to maladaptive changes in the heart, including inflammation, fibrosis, and hypertrophy ^9^. The Ang II/AT1R axis hyperactivation in pathological outcomes can be balanced by the activation of RAS protective axis ^9–14^, especially by the activation of Mas receptor (MasR) by Ang-(1-7), its endogenous ligand ^15^. Under normal conditions, the Ang-(1-7)/MasR axis activation promotes a decrease in sympathetic activity, induces vasodilation, and consequently, reduces blood pressure ^9^. The activation of Ang-(1–7)/MasR during pathological conditions exerts anti-hypertrophic, anti-inflammatory and anti-fibrotic cardioprotective effects ^14^. Although Ang-(1–7)/MasR signaling has demonstrated cardioprotective effects in several cardiovascular disorders ^16–22^, whether activation of this pathway protects the heart during TTC remains unknown.

To investigate whether activation of the Ang-(1-7)/MasR axis protects against TTC, we employed a well-established mouse model induced by a single high dose of isoproterenol (ISO) ^23–26^, which recapitulates key pathological features of the disease, including cardiomyocyte death, cardiac hypertrophy, and a robust inflammatory response characterized by the accumulation of CCR2⁺ pro-inflammatory monocytes/macrophages. Although Ang-(1-7) has been shown to promote anti-inflammatory macrophage polarization ^27^ and to exert cardioprotective effects in experimental myocardial infarction and ISO-induced cardiac hypertrophy when administered orally as a hydroxypropyl β-cyclodextrin [Ang-(1-7)-CD] ^16,17^, whether oral Ang-(1-7) attenuates the acute cardiac remodeling associated with TTC remains unknown. Therefore, we investigated whether oral administration of Ang-(1-7)-CD prevents acute cardiac hypertrophy and inflammation in a female mouse model of ISO-induced TTC.

## METHODS

### Animals

Female C57BL/6J mice aged 8-12 weeks and weighing 20-25g obtained from the Central Animal Facility at the Federal University of Minas Gerais (UFMG). Mice were housed in plastic cages (5 animals/cage) and kept under controlled temperature (24 ± 2°C) and dark-light cycles of 12h/12h, with *ad libitum* access to food and water. Environmental enrichment was provided in all cages and consisted of paper towels as nesting material and plastic igloos for shelter. All experiments were performed in compliance with the guidelines for ethical conduct in the care and use of animals and were approved by the local Ethical Committee for the use of animals (CEUA/UFMG Protocol No. 223/2023).

### Experimental groups

We used Ang-(1-7) included in hydroxypropyl β-cyclodextrin (CD) to allow its oral administration. CD increases peptide stability in gastrointestinal tract and enhances its bioavailability ^16,28^. Animals were initially allocated to experimental groups using simple randomization. To ensure baseline comparability and minimize potential bias related to body weight, all animals were subsequently weighed, and group assignments were verified to ensure that the difference in mean body weight between groups did not exceed 3 g. Female mice were randomly assigned to four groups: (i) CTRL+CD, receiving a single intraperitoneal (I.P.) injection of sterile saline (CTLR; 0.9% NaCl) and oral CD; (ii) CTRL+Ang-(1-7), receiving sterile saline I.P. and oral Ang-(1–7) included in CD [Ang-(1–7)-CD]; (iii) ISO+CD, receiving a single I.P. dose of 300 mg/Kg of (-/-) isoproterenol (ISO) and oral CD; and (iv) ISO+Ang-(1–7), receiving ISO I.P. and oral Ang-(1–7)-CD. One day prior to ISO I.P. administrations, CTRL+Ang-(1–7) and ISO+Ang-(1–7) mice began daily oral administration of Ang-(1–7)-CD (a dose of 105 μg/kg/day; 45μg/kg/day of Ang-(1-7) and 60μg/kg/day of CD) via gavage. CTRL+CD and ISO+CD groups received CD alone (60μg/kg/day). The combined Ang-(1–7)-CD formulation totaled 105 μg/kg/day (45 μg/kg Ang-(1–7) + 60 μg/kg CD). Treatment was administered daily for four consecutive days (see the experimental outline in Fig. 1A-B). The sample size was determined empirically based on previous experience with these experimental models and the reproducibility of the methods, rather than by an *a priori* statistical power calculation. At least two independent experimental batches were performed for each analysis. To minimize potential confounding factors, all animal treatments were performed at approximately the same time in the afternoon to account for circadian variation. Animals from different experimental groups were co-housed within the same cages, and cages were maintained side-by-side on the same holding racks to minimize differences in housing conditions. During sample collection and subsequent analyses, samples from all experimental groups were processed simultaneously to minimize batch effects. Euthanasia was performed by researchers blinded to group allocation.

**Figure 1:**
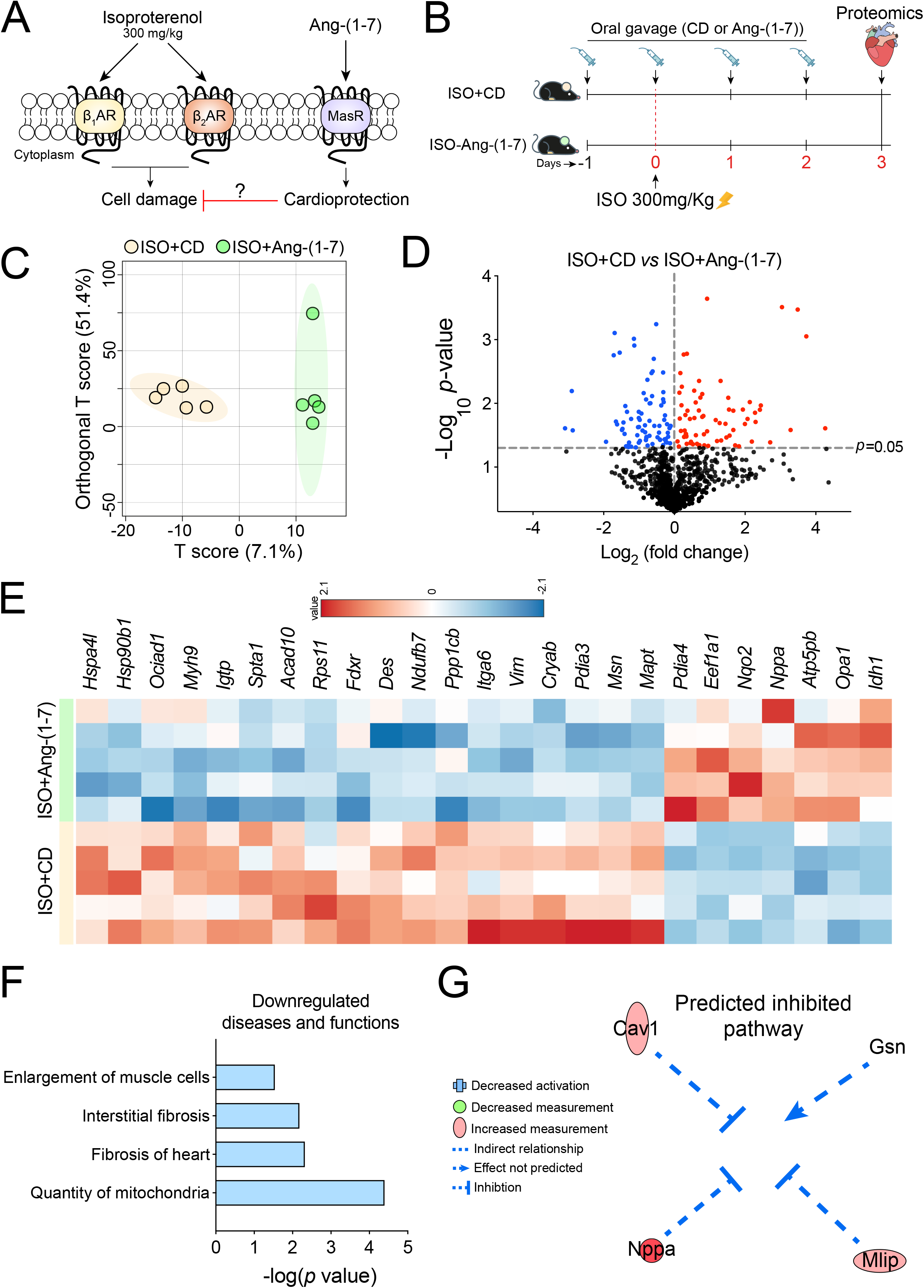
Ang-(1–7) promotes anti-hypertrophic regulation of cardiac proteins following high-dose ISO exposure. **(A)** Illustration showing how ISO interacts with both β1 and β2-adrenergic receptors resulting in cardiotoxicity, and how the Ang-(1–7)/MasR axis could exert a cytoprotective role under this context. **(B)** Experimental design depicting the oral treatment with Ang-(1–7)-CD, pharmacological induction of TTC with a single high-dose ISO exposure (300 mg/Kg), and the experimental groups distribution. At day −1, animals received the first dose of Ang-(1–7)-CD or CD alone, and treatment was given daily until day 3 following the ISO challenge. **(C)** OPLS-DA score plot showing the clear separation between experimental groups based on the proteomic signatures. Each point represents an individual sample (n=5), and the shaded areas indicate group clustering. **(D)** Volcano plot showing the differentially expressed proteins between experimental groups (*p*<0.05). Blue = downregulated and Red = upregulated. The x-axis represents the log₂ fold change, and the y-axis represents the −log10 *p*-value. Dashed lines indicate the significance thresholds (*p*=0.05). **(E)** Heatmap analysis of the differentially expressed proteins shows distinct and consistent expression patterns between experimental groups, further supporting Ang-(1–7)-driven proteomic modulation following high-dose ISO exposure. **(F)** The functional enrichment analysis on IPA shows that orally-given Ang-(1–7) induces a downregulation of pathways related to muscle cell enlargement, interstitial fibrosis, fibrosis of heart, and mitochondrial count. Bars represent −log(p-value), indicating the significance of pathway suppression. **(G)** The predicted inhibited pathway indicate reduced activation of cardiac muscle hypertrophy, driven by decreased abundance of Gsn, and increased expression of Cav1, Mlip, and Nppa, which collectively contribute to the inhibition of cardiac hypertrophy in the Ang-(1–7)-treated group.

Investigators were also blinded to group allocation during outcome assessment and data analysis for the inflammatory scoring and flow cytometry experiments.

### Pharmacological induction of Takotsubo cardiomyopathy

Takotsubo-like cardiomyopathy was induced in ISO+CD and ISO+Ang-(1–7) groups using a previously validated protocol ^24^ by a single I.P. administration of 300mg/Kg of ISO (-/-) stereoisomer (Sigma-Aldrich, Cat. No. I6504-1G) in 200 μL (sterile saline; 0.9% NaCl). The CTRL-CD and CTRL-Ang-(1–7) groups received 200 μL of sterile saline (0.9% w/v), I.P. (see experimental outline in Fig. 2A). Mice were euthanized by anesthetic overdose, receiving 80 mg/kg ketamine and 15 mg/kg xylazine intraperitoneally at day 3 after ISO or saline administration, a timepoint associated with intense inflammatory cell infiltration and a significant increase in cardiac mass ^23,24,26^. The outcome measures assessed in this study included gravimetric analysis of cardiac hypertrophy, cardiomyocyte cross-sectional area determined by wheat germ agglutinin (WGA) staining, histological inflammatory scoring, CD68 immunofluorescence, quantitative proteomics, and flow cytometry for immune-cell quantification and profiling.

**Figure 2:**
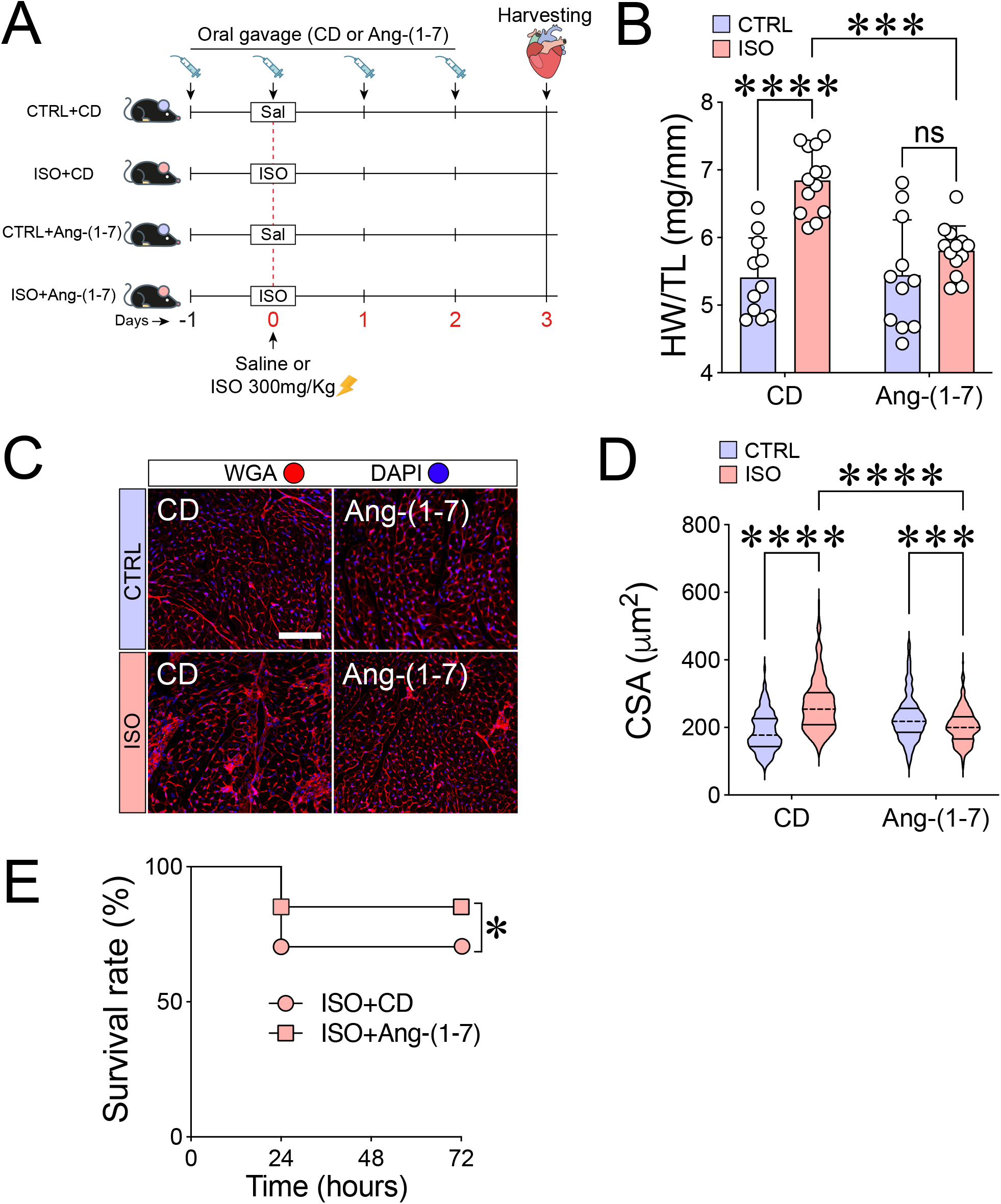
Ang-(1–7)-CD treatment prevents an increase in heart weight and an increase in the cross-sectional area of cardiomyocytes (CSA) in mice. **(A)** Overview of oral treatment with Ang-(1–7)-CD or CD alone, and TTC induction with high-dose ISO. CTRL groups received sterile saline (200 µL, I.P.) and were treated orally with Ang-(1–7)-CD or CD alone. (B) Bar graph showing that orally-given Ang-(1–7) prevented the cardiac hypertrophy induced by high-dose ISO exposure. Experimental n = hearts. CTRL+CD (n=11), ISO+CD (n=11), CTRL+Ang-(1–7) (n=13) and ISO+Ang-(1–7) (n=13). (C) Representative image of WGA-stained hearts, to evaluate cardiomyocyte cross-sectional area. **(D)** Violin plots show that Ang-(1–7) prevents cardiomyocyte hypertrophy promoted by high-dose ISO exposure. Experimental n = cells/hearts. CTRL+CD (n=56/5), ISO+CD (n=58/5), CTRL+Ang-(1–7) (n=58/5) and ISO+Ang-(1–7) (n=65/6). **(E)** Graph depicts pro-survival activity promoted by oral treatment with Ang-(1–7) following high-dose ISO exposure. Circle = ISO+CD and Square = ISO+Ang-(1–7). Experimental n = mice. ISO+CD (n=45) and ISO+Ang-(1–7) (n=40). Bar graphs show mean ± SEM. In violin plots, dashed and straight lines represent the median and the 25th and 75th percentiles, respectively. We employed a Two-Way ANOVA statistical test of variance followed by a Tukey post-hoc test (**B**), a non-parametric test (Kruskal-Wallis) followed by Dunn’s multiple comparisons post-hoc test (**D**) and a Gehan-Breslow-Wilcoxon test (**E**). In all cases the null hypothesis was rejected when *p*<0.05; *** = *p*<0.001 and **** = *p*<0.0001.

### Immunofluorescence

Mice were euthanized by anesthetic overdose and subjected to thoracotomy. A needle was inserted into the left ventricle (LV), and the hearts were perfused with 5 mL of ice-cold 1X phosphate-buffered saline (PBS). The hearts were then excised, transversely sectioned, and the basal portion fixed in 4% paraformaldehyde (PFA) for 24 hours. Samples were transferred to 30% sucrose in 1X PBS overnight. Next, the samples were embedded in optimal cutting tissue (OCT), and stored at −20°C. Cryosections of 10 µm thickness were prepared for analysis using a cryostat. To evaluate cardiomyocyte cross-sectional area (CSA) and CD68 immunolabeling, sections were washed with 1X PBS and incubated with Wheat Germ Agglutinin (WGA; Cat #W32464, Thermo) for 15 minutes. WGA binds to glycoconjugates on the cell surface, outlining plasma membranes for CSA quantification. Nonspecific binding was blocked using 3% bovine serum albumin (BSA) and 0.3% Triton X-100 in 1X PBS for 30 minutes in a humid chamber. Sections were then washed with 1% BSA and incubated with Rat anti-CD68 (1:300; Cat #MCA1957, BioRad) overnight at 4°C, followed by Alexa Fluor 488 Goat Anti-Rat (1:250; Cat #A-11006, Thermo) for 2 hours at 4°C. Nuclei were stained with DAPI for 5 minutes. The images were captured with Zeiss ApoTome 2 microscope, with a 488 and 555-nm filter. Quantification of CD68-positive macrophages and cardiomyocyte CSA was performed using Zen Blue software (RRID:SCR_013672).

### Inflammatory score

The inflammatory score was analyzed blindly in a semiquantitative manner as previously described ^23,26^. The following scores were given: 0, no infiltrate; 1, infiltrate in <25% of the ventricular myocardium; 2, infiltrates involving 25% to 50% of the myocardium; 3, infiltrates involving 50% to 75% of the myocardium, and 4, infiltrates involving 75% to 100% of the myocardium ^23^. Pictures were acquired in brightfield using a Zeiss Apotome 2 microscope at 20x magnification, and at least 20 fields were taken for each sample.

### Flow cytometry

Cardiac tissue was digested as previously described ^26,29–31^. Samples were stained with the following fluorescently conjugated antibodies (all purchased from BioLegend; 0.5 μL per 200 μL): CD45/PerCP-Cy5 (30-F11, Cat. #103132, RRID:AB_893344), Ly6G/FITC (1A8, Cat. #127606, RRID:AB_1236488), CD64/PE (X54-5/7.1, Cat. # 139304, RRID:AB_10613467), Ly6C/APC-Cy7 (HK1.4, Cat. #128026, RRID:AB_10640120), CCR2/BV421 (SA203G11, Cat. #150605, RRID:AB_2571913)and MHC-II/PE-Cy7 (M5/144.15.2, Cat. #107630,RRID:AB_2069376) for 30 minutes at 4°C and washed with FACS Buffer. The number of cells was normalized to tissue weight (cells per mg of cardiac tissue). Compensation controls were generated using UltraComp beads (Invitrogen; Catalog #01-2222-42). Data were acquired using a BD LSRFortessa flow cytometer at the Institutional Biomarkers Research Laboratory (LINBIO), College of Pharmacy, Federal University of Minas Gerais (UFMG), and analyzed using FlowJo v.X.0.7 software (RRID:SCR_008520). The gating strategy is shown in Figure 4A.

### Sample preparation for proteomics

Cell lysis was performed using 100 µL of 5% sodium deoxycholate (SDC) in 50 mM triethylammonium bicarbonate (TEAB), pH 8, 10 mM tris(2-carboxyethyl) phosphine hydrochloride (TCEP), supplemented with protease inhibitors (Complete Protease Inhibitor, Roche). To ensure complete cell lysis, samples were subjected to sonication on ice until a translucent appearance was achieved (≈15 s). Samples were then incubated for 20 minutes in a thermomixer at 45 °C to promote complete reduction of disulfide bonds and alkylation of thiol groups. Protein quantification was performed using a fluorometric assay with the Qubit system (Thermo). After quantification, the samples were diluted to achieve a final SDC concentration of 2.5%, and enzymatic digestion was carried out with trypsin (Promega) at a 1:50 enzyme-to-protein ratio for 16 h at 37 °C. The digestion was quenched by the addition of 1% formic acid. Samples were centrifuged at 14000g for 5 minutes. Digestion efficiency was assessed by mass spectrometry (Autoflex III MALDI-TOF; Bruker). Supernatant was collected, dried down using a vacuum centrifuge (speedvac; Eppendorf) and stored at −20°C until further use.

### TMT-based peptide labeling

Dried peptides were resuspended in 100 mM triethylammonium bicarbonate (TEAB) buffer at pH 8.5. Each vial of the TMT 10-plex reagent (Thermo) at room temperature was dissolved in 100% acetonitrile (ACN) and homogenized by vortexing followed by centrifugation. TMT reagent was then added to each sample, and labeling reaction was carried out for 1 hour at room temperature. To quench the reaction, 5% hydroxylamine was added, followed by incubation for 15 minutes. After labeling, samples were combined. Labelled samples were dried down using a vacuum concentrator (Eppendorf), desalted using an in-house packed (POROS R2 reversed-phase resin; PerSeptive Biosystems) microcolumn, and dried down again prior to MS analysis.

### High pH fractionation

TMT-labeled peptides were fractionated by high pH reverse-phase liquid chromatography. An UltiMate 3000 LC system (Thermo) equipped with an Acquity CSH C18 column (300 µm × 100 mm, 1.7 µm) was employed for the fractionation. Solvents for LC separation were: 20 mM ammonium formate, adjusted to pH 9.3 by NH4OH (phase A); 80% ACN, 20% 20 mM ammonium formate, adjusted to pH 9.3 by NH4OH (phase B). Peptides were eluted by a linear gradient from 2 to 50% B in 70 min collected 30 fractions were pooled into 10 fractions used for the analysis. Non-bounded peptides were collected separately and analyzed. High pH fractions were dried in a SpeedVac and stored at −20 °C until the LC-MS/MS analysis.

### Mass spectrometry analysis and protein database search

Samples were dissolved in solvent A (0.1% formic acid) and analyzed using an Orbitrap Eclipse Mass Spectrometer (Thermo) coupled to a Dionex Ultimate 3000 nanoLC system (Thermo). Samples were initially loaded onto a trap column (µ-Precolumn C18; 5 mm x 300 μm i.d.), and subsequently eluted onto an in-house packed analytical column (20 cm x 75 μm i.d., filled with Intertsil 2 µm beads). Chromatographic separation was performed using a linear reverse-phase gradient. The gradient profile was set as follows: solvent B (95% acetonitrile in 0.1% formic acid) (i) 2% to 5% over 0.5 minutes, (ii) 5% to 30% over 39.5 minutes, and (iii) 30% to 45% over 5 minutes. For washing the concentration of B has been raised to 95% over 3 minutes, followed by washing for 5 minutes, and re-equilibration with 2% solvent B. The mass spectrometer was operated in data-dependent acquisition mode (DDA) with synchronous precursor selection (SPS) and real-time search (RTS), using a 2-second cycle time. Precursor ions with charges from +2 to +6 were selected for fragmentation, and up to 10 fragments were isolated for MS3 analysis. Real-time search employed reference mouse proteome (Uniprot, obtained 2024.06.24) considering fully tryptic peptides (up to 1 missed cleavage), with TMT labeling on lysine and peptide N-terminus and carbamidomethylation of cysteine as fixed modifications, and oxidation of methionine as variable modification (up to 2 per peptide). MS1 parameters: detector – Orbitrap, resolution – 120,000 at 200 Th, mass range 350 – 1500, maximum injection time – 50 ms, AGC target – 8e4 (200%), RF lens – 30%; MS2 parameters: quadrupole isolation width – 0.7 Th, activation type – CID, normalized collision energy – 35, detector – ion trap, ion trap scan rate – turbo, maximum injection time – 35 ms, AGC target – 1e4 (100%); MS3 parameters: quadrupole isolation width – 0.7 Th, notch isolation width – 2 Th, activation type – HCD, normalized collision energy – 55, detector – Orbitrap, resolution – 30,000 at 200 Th with enhanced resolution mode, scan range – 100 – 500, maximum injection time – 54 ms, AGC target – 2.5e5 (500%). Dynamic exclusion list was set to 15 seconds with a mass tolerance of 10 ppm. Finally, peptide identification was performed against the *Mus musculus* protein database obtained from UniProt (downloaded on June 24, 2024) using MaxQuant software. Search parameters included trypsin as the specific enzyme, allowing up to one missed cleavage; fixed modifications of TMT on lysine residues and peptide N-termini; carbamidomethylation of cysteines; and variable modifications of methionine oxidation (allowing up to two modifications per peptide).

### Statistical analysis

Proteomic data was log2-transformed and normalized prior to statistical analysis. Differential protein abundance between experimental groups was assessed using unpaired two-tailed Student’s t-tests. Proteins with p<0.05 were considered statistically significant and selected for downstream analysis. Functional enrichment and pathway analyses were performed using MetaboAnalyst 5.0 (www.metaboanalyst.ca) and Ingenuity Pathway Analysis (IPA, Qiagen). On MetaboAnalyst, data were input as a protein concentration table with corresponding *p*-values, and enrichment analysis was carried out using pathway-associated metabolite sets and protein-protein interaction networks. On IPA, significantly altered proteins were mapped to curated canonical pathways and upstream regulators using the Core Analysis and TOX Analysis modules. Both tools applied over-representation analysis and right-tailed Fisher’s exact test to determine statistical significance of pathway enrichment (*p*<0.05). All reported pathways and biological functions were considered significant when p<0.05.

For functional validation data (e.g., gravimetry, cross-sectional area, inflammatory score, immunofluorescence, and flow cytometry), we employed a normality test using GraphPad Prism 10.4.0 (RRID:SCR_002798). For parametric data we used a two-way analysis of variance (ANOVA) followed by a Tukey multiple comparisons post-hoc test. For data without normal distribution, we employed a Kruskal-Wallis non-parametric statistical test followed by Dunn’s multiple comparisons post-hoc test. In both cases, the null hypothesis was rejected when the *p*-value was less than 0.05, thereby indicating statistical distinctions between the compared groups. Outliers defined as experimental points two standard deviations away from the mean were excluded. No unplanned attrition was observed during the study; any exclusions were predefined and based on technical criteria established *a priori*.

### Data availability

“The mass spectrometry proteomics data have been deposited to the ProteomeXchange Consortium via the PRIDE ^32^ partner repository with the dataset identifier PXD082872”.

<u>Reviewer access details</u>

Log in to the PRIDE website using the following details:

Project accession: PXD082872 Token: u99DPceA34xm

## RESULTS

### Ang-(1–7) reprograms the cardiac proteome signature associated with cardiac hypertrophy following a single high-dose exposure of ISO

Adrenergic discharge induced by high-dose ISO challenge leads to maladaptive changes in the heart, such as cell death, cardiac hypertrophy and inflammation, ultimately resulting in fibrosis ^23,24^. Given the well-established cardioprotective role of Ang-(1–7) ^16,17^, we sought to evaluate its protective potential in ISO-induced TTC. Mice received daily oral administration of Ang-(1–7)-CD or vehicle (CD), beginning one day prior to the single I.P. injection of ISO, and continuing for four consecutive days (Fig. 1A-B). To gain mechanistic insights into the cardiac effects of Ang-(1–7) following the adrenergic stress, we performed a TMT-based proteomics to compare the cardiac proteome signature of ISO+CD and ISO+Ang-(1–7) (see experimental outline in Fig. 1B). Multivariate analysis using Orthogonal Partial Least Squares Discriminant Analysis (OPLS-DA) revealed distinct proteome signatures between ISO+Ang-(1–7) and ISO+CD hearts, indicating different protein abundance profiles (Fig. 1C). The volcano plot analysis shown in Fig 1D depicted 150 proteins differentially regulated between ISO+Ang-(1–7) vs ISO+CD hearts (*p*<0.05). Among those, 79 were downregulated (blue dots) and 71 were upregulated (red dots). Consistently, the heatmap displayed in Fig. 1E shows the top 25 differentially regulated proteins between ISO+Ang-(1–7) and ISO+CD hearts.

We then investigated downstream molecular pathways modulated by Ang-(1–7) following high-dose ISO exposure. For this end, we used the Ingenuity Pathways Analysis (IPA) ^33^ to analyze the canonical downstream regulators potentially involved in the phenotype of ISO+Ang-(1–7) hearts. The enrichment analysis revealed several downregulated pathways related to maladaptive cardiac remodeling in ISO+Ang-(1–7) vs ISO+CD hearts, such as enlargement of muscle cells, fibrosis of heart and quantity of mitochondria (Fig. 1F). Considering that abnormal cardiac hypertrophy is a key detrimental alteration during high-dose ISO exposure, we decided to further investigate this pathway. As shown by Fig. 1G, IPA Diseases and Functions analysis predicted the inhibition of “hypertrophy of cardiac muscle” in ISO+Ang-(1–7) vs ISO+CD hearts, driven by the downregulation of gelsolin (Gsn) and upregulation of caveolin 1 (Cav1), muscular LMNA-interacting protein (Mlip) and atrial natriuretic peptide (Nppa). Collectively, these findings suggest an anti-hypertrophic role for orally administered Ang-(1–7) after high-dose ISO exposure and prompted us to examine whether the observed proteomic alterations extended to the tissue level.

### Ang-(1–7) prevents cardiac hypertrophy and promotes survival following high-dose ISO challenge

To investigate whether the oral administration of Ang-(1–7) could promote an anti-hypertrophic phenotype following high-dose ISO challenge, Ang-(1–7) were orally given starting 1 day prior to the ISO injection (see experimental outline in Fig. 2A). As depicted in Fig. 2B, high-dose ISO exposure induced cardiac hypertrophy measured by the heart-weight-to-tibial-length (HW/TL) ratio. The observed ISO-induced HW/TL increase was abolished by Ang-(1–7) treatment (Fig. 2B, *p*=0.0002 for ISO+Ang-(1–7) vs ISO+CD), showing an anti-hypertrophic activity of Ang-(1–7) in the heart exposed to the adrenergic overload. We then aimed to investigate whether this anti-hypertrophic activity could be also translated to the cellular level, by analyzing cardiomyocyte cross-sectional area (CSA). As shown in Fig. 2C–D, high-dose ISO exposure significantly increased cardiomyocyte CSA compared with control mice (ISO+CD vs CTRL+CD; *p*<0.0001), consistent with cardiomyocyte hypertrophy. Oral Ang-(1–7) significantly reduced cardiomyocyte CSA in ISO-treated mice compared with the ISO+CD group (ISO+CD vs ISO+Ang-(1–7); *p*<0.0001).

Acute exposure to high-dose ISO induces severe cardiac injury and is associated with increased mortality in mice ^24^. During the 3-day follow-up, survival was 68.4% in the ISO+CD group and 85.2% in the ISO+Ang-(1–7) group (Fig. 2E). Thus, oral Ang-(1–7) treatment increased survival by approximately 17% following ISO administration (ISO+Ang-(1–7) vs ISO+CD; *p* = 0.041). Together with the reduced heart weight and cardiomyocyte CSA, these findings show that Ang-(1–7) attenuated the hypertrophic response to high-dose ISO and was associated with improved survival.

Given the role of inflammation in cardiac remodeling following high-dose ISO exposure ^23,25,26^, we next investigated the effect of Ang-(1–7) treatment on the ISO-induced acute myocardial inflammatory response.

### Oral treatment with Ang-(1–7) abolished CCR2^+^ monocyte influx to the heart following high-dose ISO exposure

The innate immune response, particularly the recruitment of bone marrow-derived monocytes, contributes to maladaptive cardiac remodeling following high-dose ISO exposure ^23–26^. We therefore assessed the effect of Ang-(1–7) on myocardial inflammation using histology, immunofluorescence, and flow cytometry. As shown in Fig. 3A-B, ISO+Ang-(1–7) hearts had a lower myocardial inflammatory score than ISO+CD hearts (*p* = 0.043). We next assessed monocyte/macrophage accumulation by immunofluorescence staining for CD68. As expected, high-dose ISO increased the number of CD68+DAPI+ cells in the myocardium in comparison to CD-treated mice (Fig. 3C-D). Although ISO administration to Ang-(1–7)-treated mice also induced an increase in the number of CD68+ DAPI+ cells, there was a significant reduction in macrophage accumulation in Ang-(1–7) hearts compared with CD hearts in ISO-treated mice (Fig. 3C-D), supporting the hypothesis of an immunomodulatory effect of Ang-(1–7) following high-dose ISO exposure.

**Figure 3:**
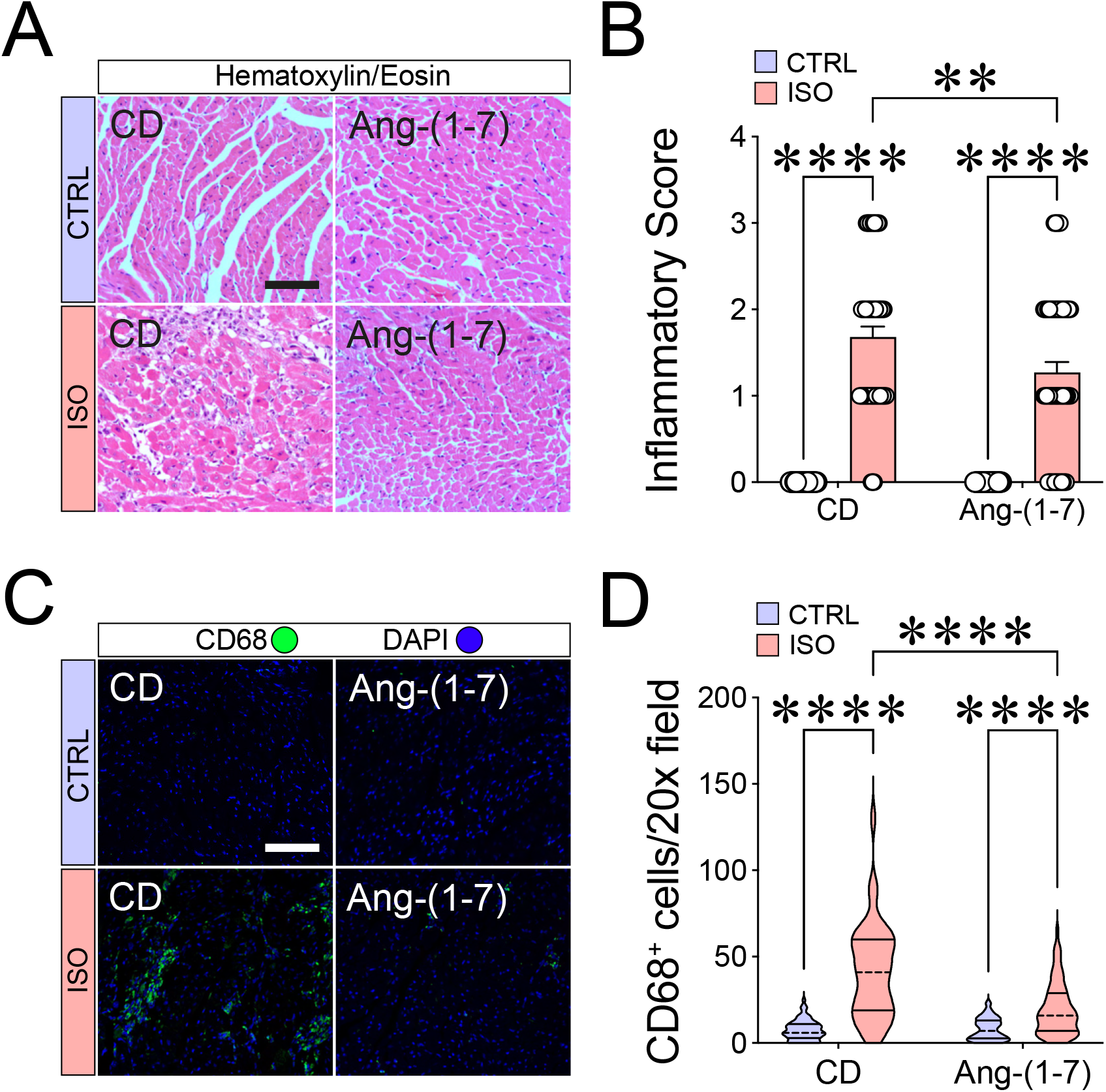
Orally given Ang-(1–7) prevents the accumulation of immune cells in the heart. **(A)** Representative images of H&E-stained hearts at day 3 post high-dose ISO exposure. Scale bar = 100 μm. **(B)** Bar graph shows that Ang-(1–7) prevents the increase of the inflammatory score elicited by high-dose ISO challenge. Inflammation scoring criteria: 0, no infiltrate; 1+, infiltrates involving <25% myocardium; 2+, infiltrates involving 25%-50%; 3+, infiltrates involving 50%-75% of the myocardium; and 4+, infiltrates involving 75%-100% of the myocardium Experimental n = fields/hearts. CTRL+CD (n=40/5), ISO+CD (n=45/5), CTRL+Ang-(1–7) (n=50/5) and ISO+Ang-(1–7) (n=59/6). **(C)** Representative images of immunostained myocardial cross-sections with CD68 antibody, to evaluate CD68^+^/DAPI^+^ cells (macrophages/monocytes) accumulation in the myocardium. Scale bar = 100 μm. **(D)** Violin plots show that oral treatment with Ang-(1–7) prevents the increase in CD68^+^ cell infiltration increased by high-dose ISO exposure. Experimental n = fields/hearts. CTRL+CD (n=68/5), ISO+CD (n=68/5), CTRL+Ang-(1–7) (n=65/5) and ISO+Ang-(1–7) (n=69/5). Bar graphs show mean ± SEM. In violin plots, dashed and straight lines represent the median and the 25th and 75th percentiles, respectively. We employed a non-parametric test (Kruskal-Wallis) followed by Dunn’s multiple comparisons post-hoc test (**B** and **D**). In all cases the null hypothesis was rejected when *p*<0.05; ** = *p*<0.01 and **** = *p*<0.0001.

To investigate this further, we characterized the leukocyte populations affected by Ang-(1– 7) following ISO-induced adrenergic stress by FAC-analyzing cardiac immune cells at day 3 (gating strategy is shown in Fig. 4A). High-dose ISO increased the cardiac abundance of CD45⁺ leukocytes (Fig. 4B), Ly6G⁺ neutrophils (Fig. 4C), and CD64^hi/int^Ly6C^hi/lo^ monocytes/macrophages (Fig. 4D) in CD-treated mice. Notably, Ang-(1-7) treatment attenuated the ISO-induced increase in these leukocyte populations, as fewer cells were observed in the hearts of Ang-(1-7)-treated mice compared with ISO-treated controls (Fig. 4B-D).

**Figure 4:**
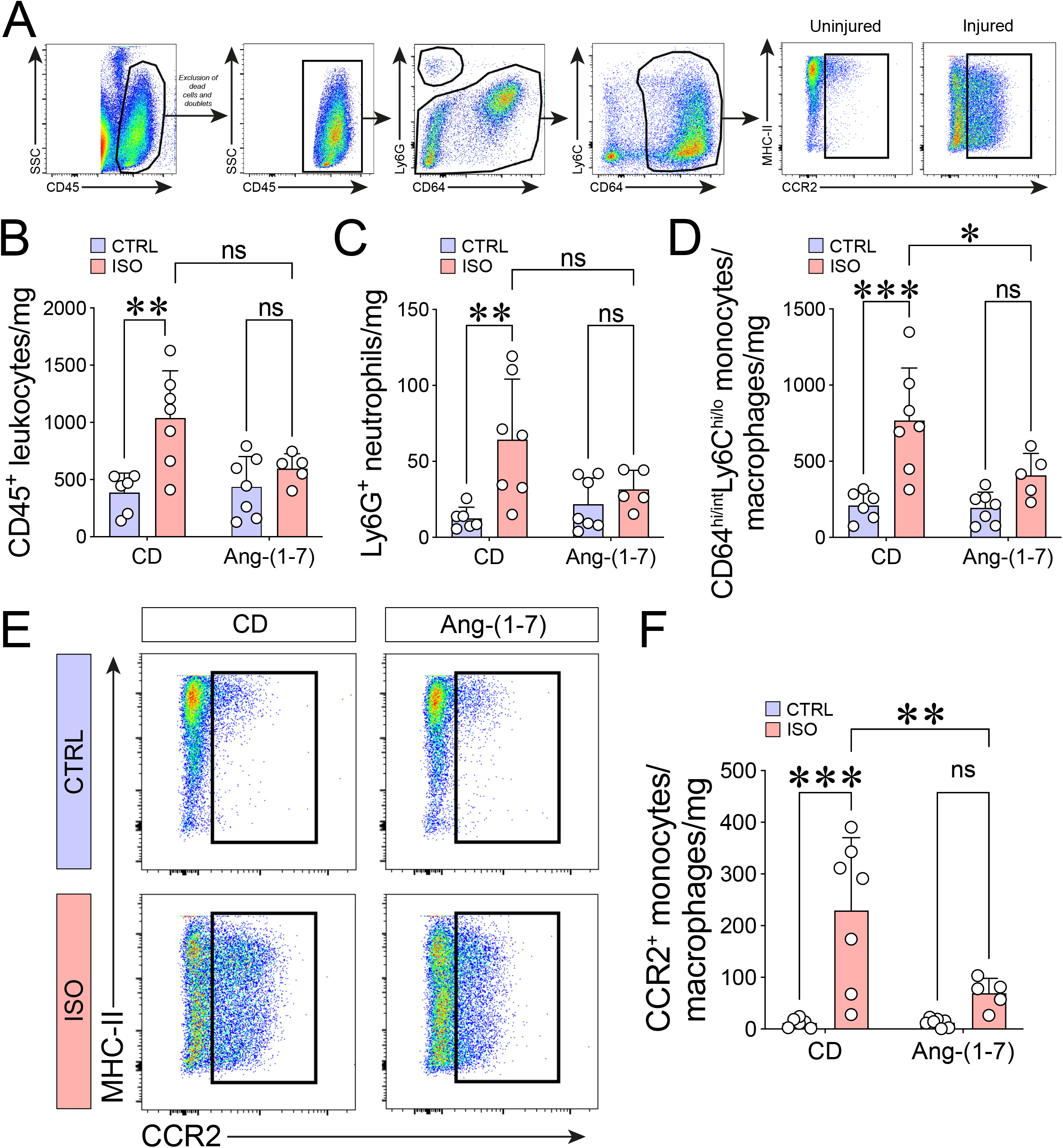
Oral treatment with Ang-(1–7) prevents CCR2^+^ monocyte influx to the heart following high-dose ISO exposure. **(A)** Gating strategy for characterization of myocardial leukocyte subpopulations using flow cytometry. **(B)** Bar graph shows that high-dose ISO challenge induces increased CD45^+^ leukocyte influx, and that treatment with Ang-(1–7) attenuates this response. Orally-given Ang-(1–7) also prevented the influx of neutrophils (Ly6G^+^ cells) **(C)**, and monocytes/macrophages (CD64^hi/int^ Ly6C^hi/lo^ cells) **(D)** that are increased by high-dose ISO exposure. **(E)** Oral treatment with Ang-(1–7) abolished the influx of CCR2^+^ cells to the heart, which is increased by high-dose ISO challenge. Experimental n = hearts. CTRL+CD (n=6), ISO+CD (n=7), CTRL+Ang-(1–7) (n=7) and ISO+Ang-(1–7) (n=5). Bar graphs show mean ± SEM. We employed a Two-Way ANOVA statistical test of variance followed by a Tukey post-hoc test (**B, D** and **F**), and a non-parametric test (Kruskal-Wallis) followed by Dunn’s multiple comparisons post-hoc test (**C**). In all cases the null hypothesis was rejected when *p*<0.05; * = *p*<0.05; ** = *p*<0.01; *** = *p*<0.001 and n.s = non significant.

Among the immune cell populations involved in acute maladaptive cardiac remodeling following high-dose ISO exposure, CCR2⁺ monocytes play an important role ^23,25,26^. Following myocardial injury, these cells are recruited from the bone marrow and differentiate into macrophages within the injured tissue, contributing to the inflammatory response ^34^ and cardiac dysfunction ^25^ in this model. High-dose ISO markedly increased the accumulation of CCR2⁺ monocytes/macrophages in ISO+CD hearts (Fig. 4E-F), whereas no significant increase was observed in mice treated with Ang-(1–7).

Overall, Ang-(1–7) reduced the acute inflammatory response to high-dose ISO, particularly the accumulation of CCR2⁺ monocytes/macrophages in the heart. Together with the attenuation of cardiac hypertrophy and improved survival, these findings support a cardioprotective effect of oral Ang-(1–7) during the acute phase of ISO-induced TTC in female mice.

## DISCUSSION

Current therapeutic approaches for TTC are primarily focused on managing associated complications, such as arrhythmias, thromboembolism, pulmonary edema, and heart failure ^4^, rather than directly protecting the myocardium from inflammation and hypertrophy. In this context, the RAS protective arm emerges as a promising therapeutic target, as Ang-(1–7) has demonstrated cardioprotective properties in other cardiac disorders ^16–22^. TTC is associated with elevated circulating Ang II levels ^6^, indicating activation of the classical RAS axis during the condition. Moreover, TTC patients receiving angiotensin-converting enzyme inhibitors (ACEi) or AT1R blockers, both of which are known to increase Ang-(1–7) levels ^35–41^, have shown improved survival rates ^42^. Although these clinical observations do not establish a direct role for Ang-(1–7) in TTC, they provide a rationale for investigating the protective RAS axis in this condition. In line with this rationale, our findings show that oral Ang-(1–7) protects against acute maladaptive cardiac remodeling in experimental TTC. Proteomic analysis showed that Ang-(1–7) modified the cardiac response to acute adrenergic stress toward an anti-hypertrophic profile. This anti-hypertrophic effect was validated at both the tissue and cellular levels, as evidenced by significant reductions in HW/TL and cardiomyocyte CSA, and was accompanied by improved survival rate. In parallel, high-dose ISO elicited a myocardial inflammation characterized by increased inflammatory score and accumulation of CD68⁺ cells and CCR2⁺ monocytes/macrophages, whereas oral Ang-(1–7) treatment attenuated these inflammatory changes, particularly CCR2⁺ monocyte/macrophage accumulation.

The anti-hypertrophic activity promoted by the Ang-(1–7)/MasR axis is well established ^14,43^ and has been demonstrated in multiple experimental models of cardiac disease ^21,22^. In the present study, we identified changes in proteins associated with cardiac hypertrophy following Ang-(1–7) treatment. Among these, muscular LMNA-interacting protein (Mlip) and natriuretic peptide A (Nppa), both implicated in limiting pathological cardiac remodeling, were increased by Ang-(1–7). Increased Mlip expression has been linked to the preservation of cardiomyocyte homeostasis and adaptive remodeling under stress conditions ^44,45^. Likewise, Nppa is a classical anti-hypertrophic mediator ^46–48^ that activates protective pathways such as cGMP/PKG signaling, thereby limiting pathological cardiac growth ^48–50^. This finding is consistent with previous studies showing that Ang-(1–7) activates NO/cGMP signaling and attenuates cardiomyocyte hypertrophy ^21^. Conversely, oral Ang-(1–7) reduced the abundance of gelsolin (Gsn), a protein implicated in cardiomyocyte growth and BNP expression ^51^, as well as in the paracrine promotion of pathological hypertrophy ^52^. Together, these proteomic changes are consistent with the attenuation of cardiac hypertrophy observed at both the tissue and cellular levels following Ang-(1–7) treatment.

Previous studies have shown that bone marrow-derived neutrophils and monocytes recruited to the injured myocardium are associated with the pathological remodeling in the TTC model ^24–26,53^. Consistent with these findings, here we observed a substantial accumulation of monocyte-derived CCR2⁺ macrophages in the myocardium at day 3 post-ISO, a cell population known to contribute to the amplification of the inflammatory response ^34^ and cardiac dysfunction in this model ^25^. Oral Ang-(1–7) prevented the ISO-induced increase in leukocytes, neutrophils, and monocytes/macrophages. Notably, the increase in CCR2^+^ monocytes/macrophages observed after ISO exposure was absent in Ang-(1–7)-treated mice.

The reduction in CCR2^+^ monocytes/macrophages is particularly relevant given the established contribution of the CCL2/CCR2 axis to pathological cardiac remodeling in several models of cardiovascular disease ^54^, including Takotsubo-like cardiomyopathy ^25^. Indeed, inhibition of the CCL2/CCR2 axis has been shown to prevent or ameliorate cardiac remodeling in models of pressure overload and preclinical HFpEF ^55,56^. Our findings are consistent with these observations and further support an association between reduced accumulation of CCR2^+^ monocytes/macrophages and the cardioprotective effects of Ang-(1-7). However, whether Ang-(1–7) acts directly on this cell population or modulates the CCL2/CCR2 axis remains to be determined. Moreover, the immunomodulatory effects of Ang-(1–7) may be secondary, at least in part, to reduced ISO-induced myocardial injury and cell death, resulting in a diminished inflammatory stimulus and consequently reduced recruitment of CCR2⁺ monocytes/macrophages.

Overall, our findings provide evidence that oral Ang-(1–7) attenuates acute cardiac remodeling in a female mouse model of ISO-induced TTC, although limitations should be considered when interpreting these findings. First, Ang-(1–7) treatment was initiated before ISO administration, allowing us to assess its protective potential but not its therapeutic efficacy when administered after TTC onset. In addition, although our proteomic and immune-cell analyses identified molecular and cellular responses associated with Ang-(1– 7) treatment, the involvement of MasR and its downstream signaling pathways was not directly investigated. Another important consideration is that the experiments were performed in young female mice with preserved ovarian function. Given that TTC predominantly affects older, postmenopausal women, the influence of estrogen on the cardioprotective effects observed here cannot be excluded. Studies using ovariectomized mice will be important to determine whether Ang-(1–7) retains its protective effects under conditions of estrogen deprivation.

## ACKNOWLEDGEMENTS

We thank Dr. Robson A. Santos for kindly providing Ang-(1–7) complexed with hydroxypropyl β-cyclodextrin (Ang-(1–7)-CD) and the corresponding vehicle (hydroxypropyl β-cyclodextrin; CD). Flow cytometry analyses were performed at the Institutional Biomarkers Research Laboratory (LINBIO), College of Pharmacy, UFMG. Generative artificial intelligence (ChatGPT, OpenAI) was used solely to assist with English-language editing and improve the clarity and readability of the manuscript. All scientific content, interpretation of the results, and final text were reviewed and approved by the authors. S.G. and TV-B are CNPq Research Fellows.

